# Collective Self-Assessment in Banded Mongoose Intergroup Contests

**DOI:** 10.1101/2025.09.23.678018

**Authors:** CW Rayner, PA Green, KL Hunt, FJ Thompson, F Mwanguhya, MA Cant, DWE Sankey

## Abstract

Contests over resources are widespread in nature. To optimize outcomes, animals assess fighting abilities, deciding to escalate conflicts based on their own strength (self-assessment) or comparing their own strength with that of their rival (mutual assessment). While most research focuses on one-on-one (dyadic) contests, the assessment strategies employed by groups remain poorly understood. Mutual assessment is frequently assumed, as more information is thought to improve decision-making; however, this assumption has rarely been tested. Here we used a dataset spanning 23 years and 641 intergroup contests in a banded mongoose (*Mungos mungo*) population in Queen Elizabeth National Park, Uganda. Our results support a model of self-assessment: groups with many males tend to escalate conflicts regardless of the rival group’s strength, thus contrasting the commonly held assumption that decisions during intergroup contests are made by mutual assessment. We suggest that assessing rival group strength during conflict could be disproportionately costly, compared with assessing own group strength, which can be done over longer time periods and is easier to obtain. Greater understanding of these dynamics can shed light on the drivers and escalation patterns of intergroup conflict across social species, including humans.

**Lay Summary:** When two rival groups come together, what determines whether or not they fight? We found support for the hypothesis that banded mongoose groups escalate into physical fights based upon an estimation of their own group’s strength, which we term collective self-assessment. We did not find evidence for a commonly held notion that groups compare their own group’s strength with their rival’s. Distinguishing between assessment strategies is intensively researched in contests between individuals but has rarely been applied to intergroup conflict. Here we applied the successful “assessment strategy framework” to group conflict, analyzing over 20 years of data from over 600 intergroup interactions. We suggest that collective self-assessment is employed because it is faster than mutual assessment, which offers an advantage in conflicts where time is of the essence.

## Introduction

Natural resources can be limited and unevenly distributed in both in time and space. This creates competition for those resources when they can be monopolized (Bornstein 2003; De Dreu *et al*. 2020; De Dreu & Triki 2022; Grover 1997; Mullon & Lehmann 2022; Nagel 1995; Rusch & Gavrilets 2020; Wrangham 1999). Contests are one way to settle disputes over such resources and are widespread across taxa (Briffa *et al*. 2013). Although the benefits from winning contests can lead to increases in fitness (Le Boeuf 1974; De Dreu & Triki 2022), the costs of escalation can be as impactful; for example, through the loss of energy (Briffa & Sneddon 2007; Hack 1997; Payne & Pagel 1996, 1997), and risk of injury (Lane & Briffa 2017; Payne 1998) or death (Enquist & Leimar 1990; Harbom *et al*. 2008; Thompson *et al*. 2017; Wrangham *et al*. 2006). As such, evolutionary theory predicts that contestants should minimize the costs and maximize the benefits when deciding whether to fight, and for how long (Birch 2016; Budaev *et al*. 2019; Parker & Smith 1990).

To respond optimally and reduce uncertainty animals can assess fighting ability (also termed resource holding potential (RHP)) (Arnott & Elwood 2008, 2009; Parker 1974; Parker & Rubenstein 1981; Smith & Parker 1976). Studies of assessment in dyadic contests have shown that individuals can assess many different RHP-contributing features of themselves and their opponents (e.g., (Arnott & Elwood 2009; Green *et al*. 2021a; Green & Patek 2018; Morrell *et al*. 2005; Pinto *et al*. 2019)), including morphology (Bohórquez-Alonso *et al*. 2014; Palaoro & Briffa 2017), physiology (Briffa & Sneddon 2007; Copeland *et al*. 2011), and behaviour (Camerlink *et al*. 2015; Wilson *et al*. 2011). This is well studied in contests between individuals (i.e., dyadic contests), but much less is known about assessment in contests between groups (but see Briffa et al. 2014).

When groups gather information to make conflict decisions, do they assess own strength (self-assessment) or do they assess both their own strength and the strength of the other group (mutual assessment)? In dyadic contests, the “assessment strategy framework” that contrasts self- and mutual assessment has been a widely utilized framework to differentiate between strategies used to assess RHP (Arnott & Elwood 2009; Fig. 1). It has been suggested that this framework can be fruitfully extended to intergroup contests (Green *et al*. 2021a). Yet, most intergroup contest studies do not properly test between assessment models. Some experimental studies that use presentations (playbacks or scent marks) have been fundamental in probing information-gathering during intergroup contests, but as no actual contest occurs during presentations it is not possible to relate them directly to how this information-gathering influences the decisions made in contests (Herbinger *et al*. 2009; Spezie *et al*. 2023; Benson-Amram *et al*. 2011;, Müller & Manser 2007; Furrer *et al*. 2011. To the best of our knowledge the assessment strategy framework has still not been appropriately implemented in an intergroup context outside of humans (Briffa, 2014). Adapting the assessment strategy framework (Arnott and Elwood, 2009) to intergroup contests can reveal potentially shared principles across levels of biological organization (individuals to groups), while suggesting how groups come to collaborative decisions during contests.

**Figure 1.**
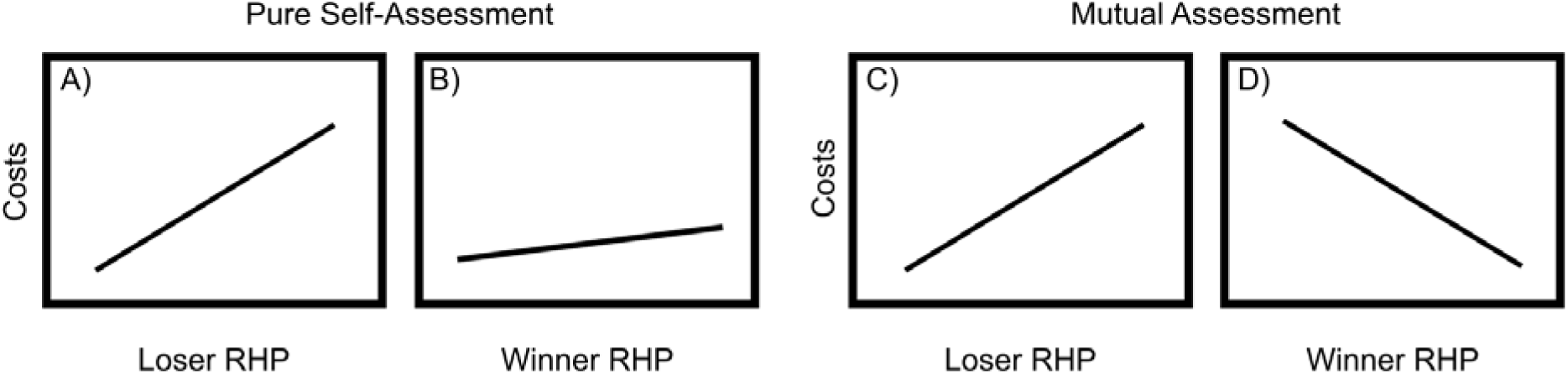
Assessment strategy framework. (Arnott & Elwood 2009). Pure self-assessment predicts **A)** a positive relationship between loser RHP and costs (e.g., duration, likelihood of escalation), and **B)** a weaker positive relationship between winner RHP and costs. Mutual assessment predicts **C)** a positive relationship between loser RHP and costs, and **D)** a negative relationship between winner RHP and costs.

The assessment strategy framework makes assumptions and predictions about the drivers of contest behaviors, which can be tested by comparing relationships between competitor RHP and contest costs (e.g., duration, likelihood of escalation; summarized in Figure 1). If animals use a pure self-assessment strategy, contests are won by those with the greater cost threshold, determined by their RHP, and no information about the opponent is assessed (Maynard Smith 1974; Patrick TREE paper). Contest costs are driven mostly by the fighting ability of losers, since winners only need to withstand a higher cost than their opponent (Fig. 1A-B) (Mesterton-Gibbons *et al*. 1996; Payne & Pagel 1996, 1997). In mutual assessment, a loser forfeits not when costs reach a certain limit, but as they discern that they have a lower RHP than their rival (Enquist *et al*. 1990; Enquist & Leimar 1983, 1987). As in self-assessment, high RHP losers can incur higher costs (Fig. 1C), yet, as opponents are assessed, high RHP winners are more quickly identified by the loser, resulting in lower overall costs of conflict for winners and losers (Fig. 1D). It is worth noting that some species can fit more than one model, depending on context, or can even change strategies during a contest (Lobregat et al. 2019; Chen et al. 2022; Dinh et al. 2020).

When applying the assessment strategy framework to the group context, we assume that group members each choose the conflict costs they are willing to incur based only on an assessment of their own group’s RHP (collective self-assessment), or the RHP of their own and the rival group’s RHP (collective mutual assessment), and that these individual choices combine to determine the decision of the group. Note as a shorthand we describe groups as taking decisions or using assessment strategies, without assuming that groups are agents in themselves (Okasha 2018).

Here, we use the assessment strategy framework to test which assessment strategy is used in escalation during intergroup contests in banded mongooses (*Mungos mungo*) – a cooperatively breeding species which hold territories and fight fiercely over resources (Cant *et al*. 2013, 2016) such as oestrus females (Green et al 2024) and food (Thompson et al 2017).. Banded mongooses are an ideal species to test for intergroup contest assessment strategy for several reasons. Firstly, banded mongoose intergroup contests are highly consequential. Intergroup conflict in this species is responsible for 10% or more of adult deaths with identifiable causes, which in mammals is matched only by chimpanzees, wolves, lions, and some human societies (Johnstone *et al*. 2020; Wrangham *et al*. 2006; Cubaynes *et al*. 2014). These high stakes likely impose strong selection pressures on assessment strategies. Second, contests have a range of escalating intensities from non-physical (e.g., war crying, forming battle lines, chasing) to physical (e.g., biting, scratching, wrestling) including injurious lethal violence (Cant *et al*. 2016) (Fig. 2). This variance in intensity (a proxy for cost) is essential for testing assessment strategy (Arnott & Elwood 2009), and similar metrics of escalation have been used in studies of dyadic assessment strategies ( Green & Patek 2018; McGinley et al 2015; Yasuda *et al*. 2012). Finally, we have known proxies for banded mongoose group RHP, which is a crucial prerequisite for testing assessment of RHP. We know that contest success is most strongly determined by the number of adult males in the group and age of the group’s oldest male (Green *et al*. 2022). Overall, the intensity and variability of conflict, and our solid baseline understanding of RHP proxies set the stage for banded mongooses to be an ideal model for testing assessment strategies in intergroup contests.

**Figure 2.**
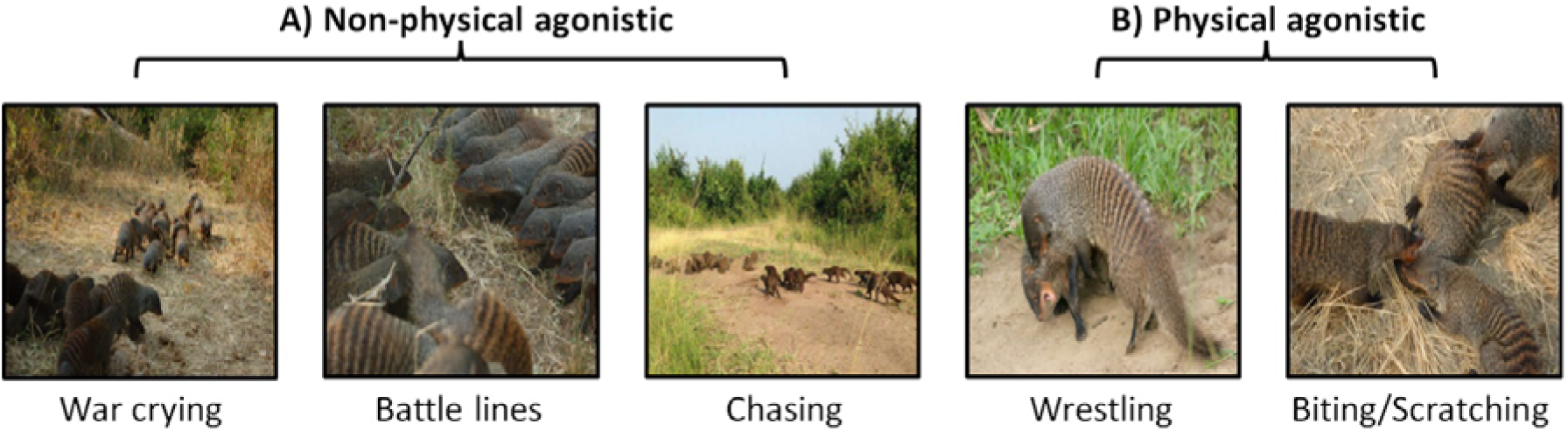
Categories of agonistic behavior observed during intergroup contests in banded mongooses. *A)* Agonistic behaviors such as war crying, forming battle lines, and chasing were considered as a “non-physical” level of intensity; ***B)*** Agonistic behaviors such as biting, scratching, wrestling, or where aggressive physical contact was observed was considered as a “physical” level of intensity. Injurious and lethal violence, where aggressive physical contact resulted in severe or fatal injury or death was also included as a “physical” level of intensity. (Image credit from left to right: Harry Marshall, Harry Marshall, Harry Marshall, Dave Seager, and Harry Marshall)

Here we tested collective self-assessment and collective mutual-assessment as two alternative hypotheses that describe patterns of escalation in banded mongoose contests. Following common assumptions in intergroup conflict, we predicted that groups would use mutual assessment. Although more cognitively demanding, the use of more information from mutual assessments can allow groups to give up earlier when clearly outmatched in fights, and is often assumed the superior or default strategy for intergroup assessment (Arnott and Elwood, 2012;Van Belle & Scarry 2015; Bennett & Stam 1996; Chan 2003; Langlois & Langlois 2009; Radford 2003; Stulp *et al*. 2012). For our measure of costs, we used the degree of escalation, from non-physical to physical (Fig. 2). If banded mongooses used collective self-assessment, we expect the previously identified RHP proxies—(i) number of adult males or (ii) the age of the oldest male (Green *et al*. 2022)—would be significantly positively correlated with the probability of escalation to physical violence for losing groups and would show a weaker positive relationship in winning groups (Fig. 1A-B). Conversely, if, as predicted, banded mongooses do employ collective mutual assessment, we expect these same variables to exhibit a significant positive correlation in losing groups and a significant negative correlation in winning groups (Fig. 1C-D).

## Methods

### Study Population and Data Collection

All data was collected as part of the Banded Mongoose Research Project, a long-term study on a population of banded mongooses in and around Mweya Peninsula, Queen Elizabeth National Park, Uganda (0°12′S, 29°54′E). This study includes data collected from 2^nd^ February 2000 to 7^th^ February 2023 encompassing 641 intergroup contests from 43 different groups and 81 unique pairings of groups. In general, at any given time, there are approximately 250 individuals present within the population making up 10-12 groups consisting of around 10-30 adults each (Cant 2000; Cant *et al*. 2013). Every 1-3 days researchers recorded data on life-history (e.g., births, deaths), composition of each group, and details about their intergroup interactions (below), among other records not relevant to the present work.

### Scoring Intergroup Contest Intensity and Escalation

Intergroup contests were recorded opportunistically as they occurred. Intergroup contests were defined as any time when at least two groups directed agonistic behavior towards each other (Fig. 2). Intensity and escalation of intergroup contests were scored using comments recorded by the Banded Mongoose Research Project team. As an intergroup contest often involves a series of behaviors in which intensity escalates until one or both groups retreat (at which point the intergroup contest ends) the highest point of escalation was used as the level of intensity for each contest. For ease of analysis, the range of intensities possible during an intergroup contest was divided into two categories (Fig. 2). The lowest level of intensity was “non-physical” which was defined as an instance where two groups directed non-physical agonistic behavior (e.g., vocal and visual displays; (Cant *et al*. 2016)) towards each other and/or fled upon sighting. If this then escalated to fighting between the two groups in which there was aggressive physical contact (e.g., biting, scratching, wrestling; (Cant *et al*. 2016)), this was defined as “physical”. Intergroup contests which escalate to physical combat are expected to have higher contest costs (such as energy used and injury risk) compared to non-physical contests (Green & Patek 2018; Lane & Briffa 2017; McGinley *et al*. 2015). Our scoring approach was then translated into a binomial escalation metric with 0 representing non-physical and 1 representing physical. Injurious and lethal violence in which aggressive physical contact results in severe injury or death of one or more individuals was included in the “physical” category, rather than a category of its own. This is because, firstly, injury and mortality were data poor (9.4% of all intergroup interactions; N = 60), but more importantly, whether individuals suffer an injury is not so much a decision as it is a (potentially random) outcome of such physical combat. By contrast, whether violence escalates from non-physical to physical combat is a decision the group may make. Therefore, our use of the escalation metric likely reflects the cost each group was willing to pay.

For each intergroup contest, a qualitative comment (a description of the events of the contest) was recorded by observers in the field. These comments were then assessed by three researchers (CR, FM, DS) to evaluate whether the contest escalated into physical violence (with 0 representing non-physical and 1 representing physical), or whether the comments did not allow us to determine whether or not the contest escalated (termed: “undeterminable”). Second, all researchers assigned a confidence score (1–3) to each of their categorizations of intensity score, reflecting their certainty in the intensity score. The confidence score was based on criteria such as the clarity and detail of the recorded comment, as well as contextual factors that might influence interpretation (e.g., visibility conditions or proximity to the event). A score of 1 indicated high confidence, 2 indicated moderate confidence, and 3 indicated low confidence. We ended up removing contests that scored with low confidence (3) from the dataset because they were missing data on other variables included as a fixed effect in our analysis. The three researchers discussed any ambiguous comments in detail to assign an escalation and confidence score if possible. The discussion was guided by FM, who has over 27 years of experience observing and collecting field data on banded mongooses, and managing the Banded Mongoose Research Project .

We report models using both highly confident and moderately confident scores (1,2), and models including only highly confident scores (1).

### Statistical Analysis

All statistical analyses were carried out using *R* 4.01 (Team 2021). Binomial escalation response data were analyzed using generalized linear mixed effect models (GLMMs) with a binomial error structure and a logit link function (Bolker *et al*. 2009) with the ‘lme4’ package (Bates *et al*. 2015). For both losing groups and winning groups, we tested the effect of two RHP predictor variables— number of adult males (>6 months old) and age of the oldest male (days) (Green et al. 2022). Both variables were scaled using the scale function (Becker *et al*. 1988).

In total, we ran two GLMM models, one model using data from high and moderate confidence scores, and another using data from only high confidence scores. The model for each confidence category is as follows (“∼” represents “as predicted by”): 1) escalation ∼ loser number of adult males+ loser age of oldest male+ winner number of adult males+ winner age of oldest male. Models without interactions between these variables were simpler and a better fit than models including interactions. Winner group ID, loser group ID, and unique pairings of groups (winner group ID + loser group ID) were included as random intercepts in every model to account for repeated measures of intergroup contests between the same groups and group dyads. Test statistics were obtained using the Anova function and confidence intervals using bootstrapping. We present the p-values, chi-squared values , parameter estimates (β) on the logit-scale, standard errors, and confidence intervals of each GLMM model, and compare p-values and direction of parameter estimates to the assumptions of pure self-assessment, and mutual assessment (Fig. 1). The collective self-assessment hypothesis would be supported if there was a positive and statistically significant relationship between RHP predictor variables and escalation for losing groups (β>0; p<0.05) and a weaker relationship for winning groups (no significance threshold) (Fig. 1A-B). The collective mutual assessment hypothesis would be supported if these same tests indicated a significant positive relationship for losing groups (β>0; p<0.05) and a significant negative relationship for winning groups (β<0; p<0.05) (Fig. 1C-D).

## Results

Overall, of the 641 intergroup contests we observed, 280 were ‘non-physical’ (43.7%), 230 were ‘physical’ (35.9%), and 131 were undeterminable (20.4%).

Under the analysis where high and moderate confidence scores of our escalation metrics were combined (N = 253), our results supported self-assessment: there was a significant positive relationship between the probability of escalation and number of adult males for groups that lost fights (losers; β = 0.47, SE = 0.18, χ^2^ = 7.13, 95% CI [0.14, 0.86], p=0.008), and a non-significant, positive relationship between these variables for groups that won fights (winners; β = 0.21, SE = 0.17, χ^2^ = 1.61, 95% CI [-0.13, 0.57], p=0.20) (Fig. 3A-B; Table 1), supporting the hypothesis. Support remained for the self-assessment hypothesis when using only high confidence score escalation metrics (N = 229). Here, loser groups still showed a greater probability of escalation with larger numbers of adult males (losers; β = 0.52, SE = 0.19, χ^2^ = 7.15, 95% CI [0.14, 0.95], p=0.008), with a non-significant, positive relationship between escalation and number of adult males for winners (winners; β = 0.31, SE = 0.18, χ2 = 2.82, 95% CI [-0.05, 0.71], p=0.09; Table 1). This is as expected under self-assessment.

**Figure 3.**
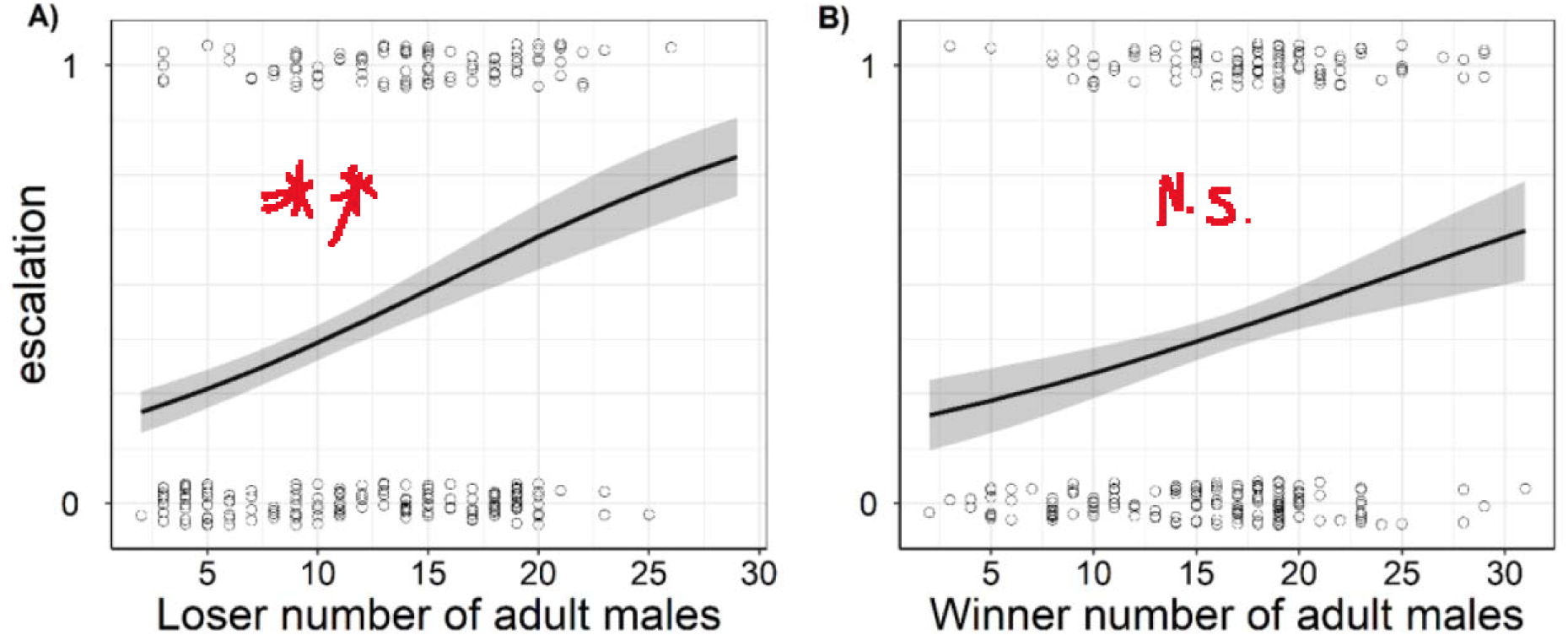
Support for collective self-assessment of number of adult males in banded mongooses. Probability of escalation plotted against **A)** loser number of adult males (p=0.008); **B)** winner number of adult males (p=0.20). (Escalation was a binary metric where contests were either 0 = non-physical, and 1 = physical). The grey shaded area shows the standard error around the fitted line. Data (circles) represent individual contests and is randomly jittered on the y-axis. Data in plot includes both high and moderate confidence scores of our escalation metrics (N = 253))

**Table 1.**
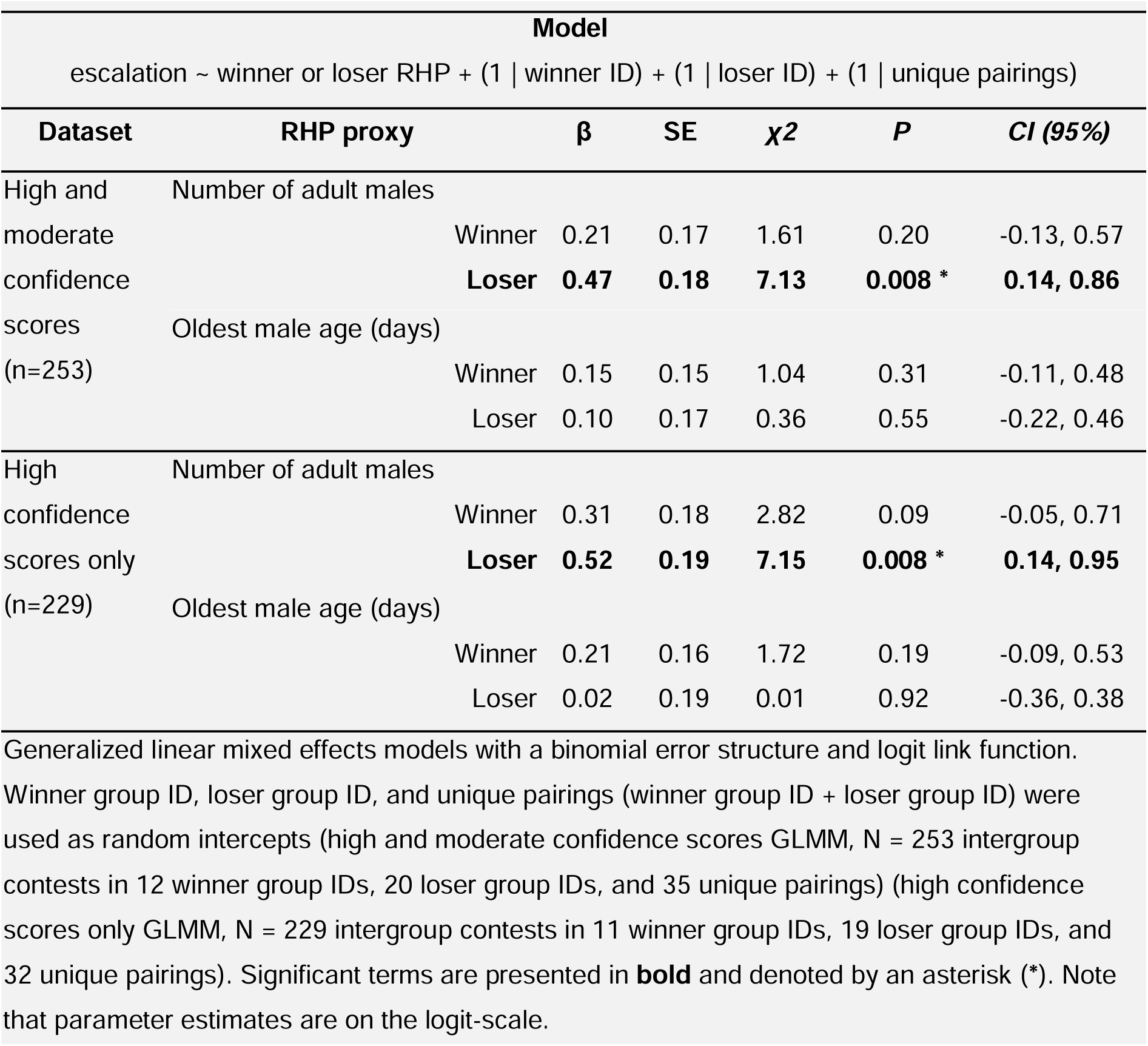
Models predicting the probability of escalation for loser or winner RHP in intergroup contests. The estimate (β), standard error (SE), chi-squared value (χ2), p-value (P), and confidence interval (CI) of a model predicting the escalation of loser or winner RHP in intergroup contests.

There was no significant relationship between oldest male age and the probability of escalation for winners or losers in either confidence category, providing no statistical support for either model of assessment (Table 1).

## Discussion

We found support for the hypothesis that mongoose groups used collective self-assessment when deciding to escalate a contest into physical violence. Specifically, our results suggest that losing banded mongoose groups were more likely to escalate in contests when they had many adult males (a known proxy of RHP; Green et al. 2022), irrespective of the number of adult males in the winning group. By contrast, we found no support for any assessment model based on the oldest male age within the group. Despite assumptions that intergroup contests are determined through mutual assessment (e.g., (Van Belle & Scarry 2015; Bennett & Stam 1996; Chan 2003; Langlois & Langlois 2009; Radford 2003; Stulp *et al*. 2012)) we found no support for this in our study.

At first sight our results are surprising. Why should banded mongooses assess only their own fighting force, and ignore potential valuable information about the outgroup? Research in dyadic contests has suggested that, while mutual assessment can minimize the costs of conflict by allowing the weaker party to back down earlier, acquiring information about an opponent may be constrained or entail prohibitively high costs (Guillermo-Ferreira *et al*. 2015; Mesterton-Gibbons & Heap 2014). For instance, assessing the strength of outgroups in the short time-window before conflict could be disproportionately costly, compared with own-group assessment which can be attained at relatively low cost over longer time periods such as through play-fighting and intragroup contests (Nolfo *et al*. 2021; Turner *et al*. 2020; Weller *et al*. 2020).

Our finding that the number of adult males in a banded mongoose group is associated with the probability of escalation accords with previous evidence of the importance of adult males for intergroup fighting success (Green *et al*. 2022). Males suffer greater mortality from conflict than females (Johnstone *et al*. 2020) and there is experimental evidence that in simulated encounters males were more likely to approach caged intruders (Cant 2002). This suggests that males are the main participants in conflicts. It has also been suggested that males have evolved adaptations as a result, such as greater body mass and head size (Green *et al*. 2022), that may be important for fighting behavior (Briffa *et al*. 2013; Chelliah & Sukumar 2013; Gvoždík & Van Damme 2003; Jennings & Gammell 2013; Vieira & Peixoto 2013; Wroe *et al*. 2005) and determining contest success (Husak *et al*. 2006; Huyghe *et al*. 2005; McLean & Stuart-Fox 2015). More closely-related to intergroup conflict, our findings are consistent with the male warrior hypothesis, which proposes that male mammals possess specific adaptations for intergroup fighting and participate more in intergroup conflict than females, and thus have a pronounced effect on group RHP (Muñoz-Reyes *et al*. 2020; Smith *et al*. 2022). Therefore, the number of adult males is likely an important and informative RHP proxy which can be assessed when making decisions during intergroup contests in banded mongooses.

Additionally, numerical superiority appears to play such an important role in intergroup fighting because several attackers can completely overwhelm a lone opponent. In banded mongooses most fatalities seem to occur when individuals become isolated and overwhelmed by groups of attackers (*personal observations*; e.g., Video S1). Outnumbering one’s opponent presents many advantages such as the ability to attack simultaneously from multiple angles, to pin down an opponent while others deliver bites or blows, or to take turns in energetically intense fighting (Adams & Mesterton-Gibbons 2003; Baglione *et al*. 2010; Nunn & Deaner 2004; Rusch & Gavrilets 2020; Wilson *et al*. 2002). The importance of numerical superiority and concentration of force is also emphasized in models of warfare between human groups (Lanchester 1916; Rapoport & von Clausewitz 1968). As discussed above, assessing own-group number of adult males may be a low-cost, more accurate way to use this information in competitive decision-making.

Despite the importance of oldest male age for banded mongoose group RHP (Green *et al*. 2022), the age of the oldest male in the group was not associated with escalation or show support for any assessment strategy. While age may play a role during intergroup contests and assessment in general (Briffa & Lane 2017; Fawcett & Johnstone 2010; Green *et al*. 2022), number of adult males may be a faster, simpler, or more reliable cue on which to base escalation decisions in intergroup contests. Assessing several attributes for more than one individual may be a cognitively demanding process (Budaev *et al*. 2019; Johnson & Fowler 2013; Tecwyn *et al*. 2017). Age in general may also be less conspicuous (Elwood & Arnott 2012; Fawcett & Mowles 2013; Nieder 2020), scalable or countable (Akre & Johnsen 2014; Bonanni *et al*. 2011; Nieder 2020) making estimates of age more prone to error. Additionally, in banded mongooses, the age of the oldest male was a weaker predictor of contest outcomes compared to number of adult males (Green *et al*. 2022), which is consistent with its weakness in association with contest escalation here.

Despite finding support for self-assessment in this analysis, intergroup conflict in banded mongooses does not fit classical assumptions of pure self-assessment models. The original self-assessment models assume no physical contact and that the costs of conflict are only gradual escalation of display intensity over time (Bishop & Cannings 1978). In banded mongooses, frequent physical aggression and severe costs (e.g., injuries, deaths) violate these assumptions (Cant *et al*. 2013; Thompson *et al*. 2017). This divergence mirrors findings in the dyadic contest literature, where violations of classical assumptions are widespread. For instance, Pinto et al. (2019) report that in 34 of 36 species in which self-assessment was supported, assumptions such as no physical contact and single-phase (display ritual) contests were violated. Many dyadic contest studies use a range of different proxies for cost, such as latency to approach and distance travelled (e.g., (Beeching 1992; Wilson *et al*. 2011)), action rate (e.g., (Hack 1997; Jennings *et al*. 2012; Pratt *et al*. 2003)), and escalation (Green & Patek 2018; McGinley *et al*. 2015; Yasuda *et al*. 2012). Subsequently, and mirroring calls in dyadic research (Pinto et al 2019), we advocate for further theoretical development that builds additional realism into the study of assessment strategies (Kokko 2013; Mesterton-Gibbons & Heap 2013).

Moving forward, key differences between individual and group contests may demand a more dedicated intergroup-specific theoretical framework. Not only do intergroup conflicts involve collective decision-making (Sankey et al., 2022), and variation in leadership dynamics (Hunt et al., 2024), but it is also possible that contests between groups could escalate into violence through mechanisms other than assessment-based decisions. For example, collective decisions to escalate into conflict may be dependent on the presence of a key individual (with an inherently risk-prone personality) already committed to escalation (Glowacki & McDermott, 2022). No collective assessment is necessary, just a propensity for the group to follow those individuals which commit themselves to conflict initiation. (The incentive for followers to join is thought to increase because key individuals pay a larger share of the startup costs – De Dreu *et al*. 2016; Gavrilets and Fortunato, 2014). We modelled the dynamics of this alternative hypothesis but found continued support for collective self-assessment in banded mongooses (See Supplemental Material). In future work, expanding and formalizing intergroup-specific contest models will help determine the generality of various intergroup assessment (or non-assessment-based) strategies across taxa.

## Acknowledgments

We would like to thank all of Michael Cant’s Socialis lab group for helpful meetings and discussion. We thank the whole Banded Mongoose Research Project team for all their support, data collection, guidance, and insight which has been invaluable throughout. In particular, we thank the Uganda Wildlife Authority and the Uganda National Council for Science and Technology for permission to carry out our research; the Wardens of Queen Elizabeth National Park for logistical support; and Solomon Kyabulima, Kenneth Mwesige, Robert Businge, and Solomon Ahabyona for helping collect data in the field. We are grateful to Harry Marshall and Emma Vitikainen for curation and maintenance of the long-term data, and Jason Gilchrist, Sarah Hodge, Matthew Bell, Corsin Müller, Neil Jordan, Bonnie Metherell, Roman Furrer, David Jansen, Jenni Sanderson, and Beth Preston for valuable contributions to the project.

